# Mathematical model of a cytokine storm

**DOI:** 10.1101/2022.02.15.480585

**Authors:** Irina Kareva, Faina Berezovskaya, Georgy Karev

**Affiliations:** Computational and Modeling Sciences Center, Arizona State University, Tempe, AZ, 85287, USA; Department of Mathematics, Howard University, Washington, DC, 20059; National Center for Biotechnology Information, National Institutes of Health - Bldg. 38A, 8600 Rockville Pike, Bethesda, MD 20894, USA.

**Keywords:** cytokine release syndrome, CRS, cytokine storm, mathematical model, IFN-gamma, IL-6, second touch hypotheses

## Abstract

Cytokine storm is a life-threatening inflammatory response that is characterized by hyperactivation of the immune system, and which can be caused by various therapies, auto-immune conditions, or pathogens, such as respiratory syndrome coronavirus 2 (SARS-CoV-2), which causes coronavirus disease COVID-19. While initial causes of cytokine storms can vary, late-stage clinical manifestations of cytokine storm converge and often overlap, and therefore a better understanding of how normal immune response turns pathological is warranted. Here we propose a theoretical framework, where cytokine storm phenomenology is captured using a conceptual mathematical model, where cytokines can both activate and regulate the immune system. We simulate normal immune response to infection, and through variation of system parameters identify conditions where, within the frameworks of this model, cytokine storm can arise. We demonstrate that cytokine storm is a transitional regime, and identify three main factors that must converge to result in storm-like dynamics, two of which represent individual-specific characteristics, thereby providing a possible explanation for why some people develop CRS, while others may not. We also discuss possible ecological insights into cytokine-immune interactions and provide mathematical analysis for the underlying regimes. We conclude with a discussion of how results of this analysis can be used in future research.

## Introduction

Cytokine storm, a life-threatening inflammatory response involving elevated levels of cytokines and hyper activation of the immune system, has recently gained particular note as one of the causes of morbidity and mortality from coronavirus disease COVID-19 (1). It has previously been observed in a variety of other circumstances, including graft vs host disease (2) and other viral infections, such as SARS (3); cytokine storms have also been implicated as one of the key culprits in the severity of the 1918 Spanish flu pandemic (4). Additionally, cytokine storms have been observed as a side effect of certain anti-cancer therapeutic interventions, such as chimeric antigen receptor, of CAR-T cell therapy (5) and bispecific T cell engagers, also known as BiTEs (6). One of the most notable therapy-induced instances of cytokine storm was the case of a Phase I clinical trial of monoclonal antibody TGN1412, which resulted in severe damage to the health of six volunteers that participated in the trial despite very accurately chosen initial doses that were administered to them (7); numerous additional reports of the details of the case can be found in the literature.

Cytokine storms are most often characterized by severe lung infections, which can lead to respiratory distress, multi-organ failure, sepsis and in some cases, death (5,8,9). Mechanistically, cytokine storms are mitigated by cytokines, which are molecules involved in supporting and regulating the immune response. Cytokine interactions form complex networks, geared towards mounting fast and efficient immune response against pathogens while also preventing excessive damage to normal tissues. If these interactions become destabilized, cytokine storms, or hypercytokinemia, may occur, where immune response causes greater collateral harm than benefit. Some prominent cytokines that are elevated during cytokine storms include interferon (IFN)-gamma, tumor necrosis factor (TNF)-alpha, as well as interleukins (IL)-6,8 and 10 (1,3,5,8,9). More generally, cytokine storms appear to reflect a scenario when the response to a pathogen, or an immune stimulatory agent, rather than a pathogen itself, results in pathology, and this is the mechanism that we wish to explore in greater detail.

Notably, while they are often used interchangeably, there exists a distinction between the terms “cytokine storm” and “cytokine release syndrome” (CRS). Cytokine storm typically refers to an acute reaction, while CRS typically refers to a more delayed response. There exists a discussion about qualitative differences between the two responses, how they are triggered and how they proceed (5), although it appears that the final qualitative dynamics are very similar between the two. Henceforth we will be using the term cytokine storm; however, we believe that the proposed model can be used for better understanding of CRS as well.

Several mathematical models have been developed to try to create and formalize a framework for better mechanistic understanding of cytokine storm dynamics. Waito et al. (10) proposed a mathematical model of cytokine storm, where they grouped cytokines into 7 categories based on their pro- and anti-inflammatory properties. They use the model, parameterized with mouse data, to describe the mutual influence of cytokine groups on each other during a cytokine storm. Yiu et al. (11) developed a large scale eighteen-order mathematical model to analyze the data from the TGN1412 clinical trial, using principal component analysis to reveal functional cytokine clusters that were specific to this case. Hopkins et al. (12) created a model of 9 major cytokines affecting the outcome of CAR-T cell based therapy. A smaller more conceptual model was proposed by Baker et al. (13), where a two-dimensional system of equations captured interactions between pro- and anti-inflammatory cytokines, displaying large regions of bi-stability and oscillations reminiscent of immune behavior in rheumatoid arthritics; the model was later extended by other authors, such as by Zhang et al. (14).

Here we propose a conceptual mathematical model that is aimed to capture general phenomenology of transition from norm to storm rather than the intricate details of cytokine biology and interactions. We use the model to identify within a theoretical framework what factors may be critical to result in this transition. The model is coupled with a model of viral infection to initiate the immune-cytokine dynamics, which can be substituted with a different sub-model depending on the question, since, according to (8), although the initial drivers leading to cytokine storm dynamics may differ, late-stage clinical manifestations of cytokine storm converge and often overlap, and therefore we expect the proposed modeling framework to be translatable for different causes.

Through our analysis, we identify key processes that within this framework can result in storm-like behavior. We demonstrate existence of a sequence of regimes as one transitions from normal to storm-like behavior, that is parameter dependent. We show the impact of both intrinsic individual-specific characteristics and infection-specific characteristics that need to converge in order to result in a cytokine storm. We analyze the immune-cytokine dynamics from an ecological point of view, showing that their interactions can shift from stabilizing predator-prey like dynamics to mutually augmenting mutualistic relationship, and show how these shifts are reflected in normal vs pathological dynamical behaviors. Finally, we show that the proposed model predicts existence of “long-haulers”, patients with chronic persistent infections, which have been observed in COVID-19, and that it predicts infection-induced autoimmunity. We conclude with a discussion of next steps and potential experiments to be designed to test predictions generated by this model to potentially identify patients that may be at a higher risk of developing a cytokine storm.

## Model Description

The proposed model consists of two subsystems: immune-cytokine subsystem (primary), and an SIV (susceptible-infected-virus) sub-system (secondary) that serves to provide sufficient perturbation to the immune-cytokine system to initiate an immune response.

Even before running simulations, we would expect to see the following types of responses:

1. Normal response: after external perturbation to the immune system subsides (infection is cleared), immune-cytokine populations return to pre-infection equilibrium.
2. CRS: even though external perturbation to the immune system has subsided (infection has been cleared), immune cells and cytokines continue affecting each other even in the absence of external stimulus.

Notably, the goal of this work is to describe a mathematical model that can capture and reproduce these behaviors, and to analyze conditions for when one or the other type of behavior will occur.

### Viral subsystem

In order to describe the impact of a viral infection on the immune system, we adapt an SIV model described in (15). We consider the dynamics of the following 3 variables: susceptible cells S(t), infected cells I(t) and viral particles V(t). We assume that the population of susceptible cells S(t) undergoes normal turnover described by *S_in_* – *k_S_S*(*t*), and can be infected by the virus at a rate b, creating infected cells I(t). Infected cells can die at a rate *k_I_* or can be cleared by immune cells *x*(*t*) at a rate *γ*. Viral particles V(t) are produced by the infected cells I(t) at a rate *v_in_* and get cleared at a rate *k_v_*. These mechanisms are described by system (1)

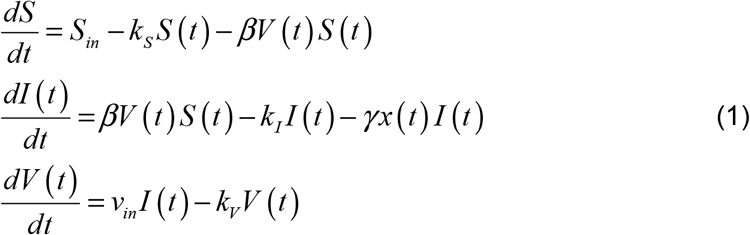

This proposed model is of course highly simplified and primarily serves the purpose of introducing a dynamic perturbation to the immune-cytokine subsystem; as such, it will not be fully analyzed. It is used here instead of a simple mechanical perturbation to the immune-cytokine subsystem to allow us to describe a variety of situations, such as chronic infection. It can be modified and adapted to different questions as needed.

### Immune-cytokine subsystem

The following system of equations aims to capture the qualitative aspects of the dynamical relationship between immune cells x(t), and two types of cytokines y(t) and z(t) that can regulate immune activity and that appear to act synergistically in hyperactive immune response (16).

First, we describe the dynamics of y(t), which are involved in direct regulation of T cells; these can be interpreted as TNF-alpha or IFN-gamma. We also describe the dynamics of z(t), which can stimulate production of y(t) and thus indirectly regulate immune cells x(t); these species can be interpreted as interleukins, such as IL-6.

We assume that cytokines y(t) have a normal turnover rate and thus maintain an infection-free baseline level 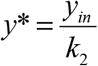. We assume that interleukins z(t) are produced in response to interactions between immune cells x(t) and cytokines y(t), and are cleared at a natural rate *a*_2_. Finally, the dynamics of immune cells x(t) is described as follows: we assume that immune cells have a normal turnover rate to maintain a normal infection-free level *x*\*. Immune cell population can additionally increase in response to infection, as captured by the term *γx*(*t*)*I*(*t*). Finally, we assume that there exists a threshold *m*, beyond which immune cells receive an additional growth boost; we interpret the existence of threshold m to be with in accordance with the second touch hypothesis (17), where antigen-experienced T cells require a “second touch” by the necessary antigen to achieve full immune activation, resulting in part in a delay between antigen encounter and immune cell expansion. The duration of additional immune cell expansion is regulated by cytokines y(t) as follows: we assume that there exists a range of concentrations of y(t) that acts as immune stimulatory, and a concentration that can become immune inhibitory. We assume that the immune cells have an additional positive growth term when concentration of cytokines is between y_1_<y(t)<y_2_, thereby capturing in a phenomenological way the dual regulatory and inhibitory property of cytokines on the immune system.

The resulting system then takes the following form:

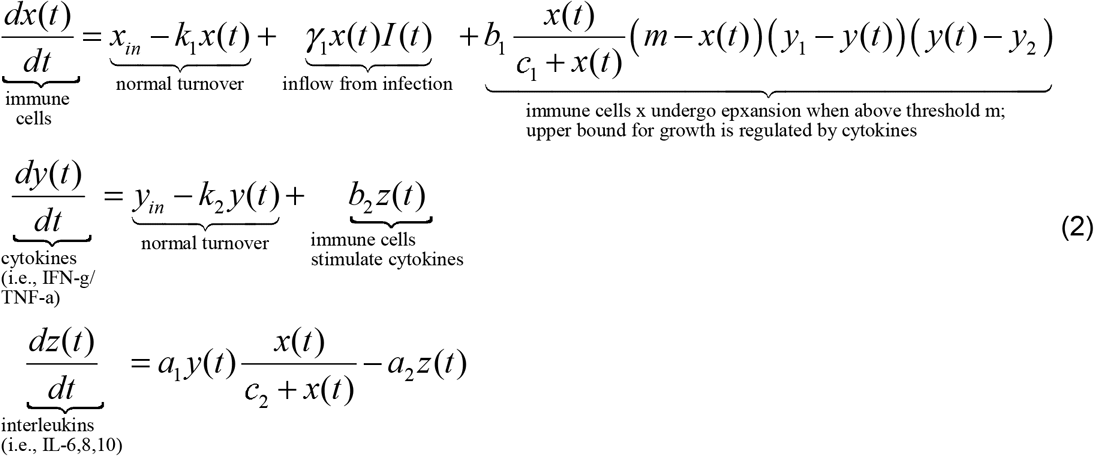

Schematic representation of this model structure is given in Figure 1A. Notably, disease-free equilibrium has to satisfy *x*\* < *m*, which is necessary to capture antigen-induced immune cell expansion.

**Figure 1.**
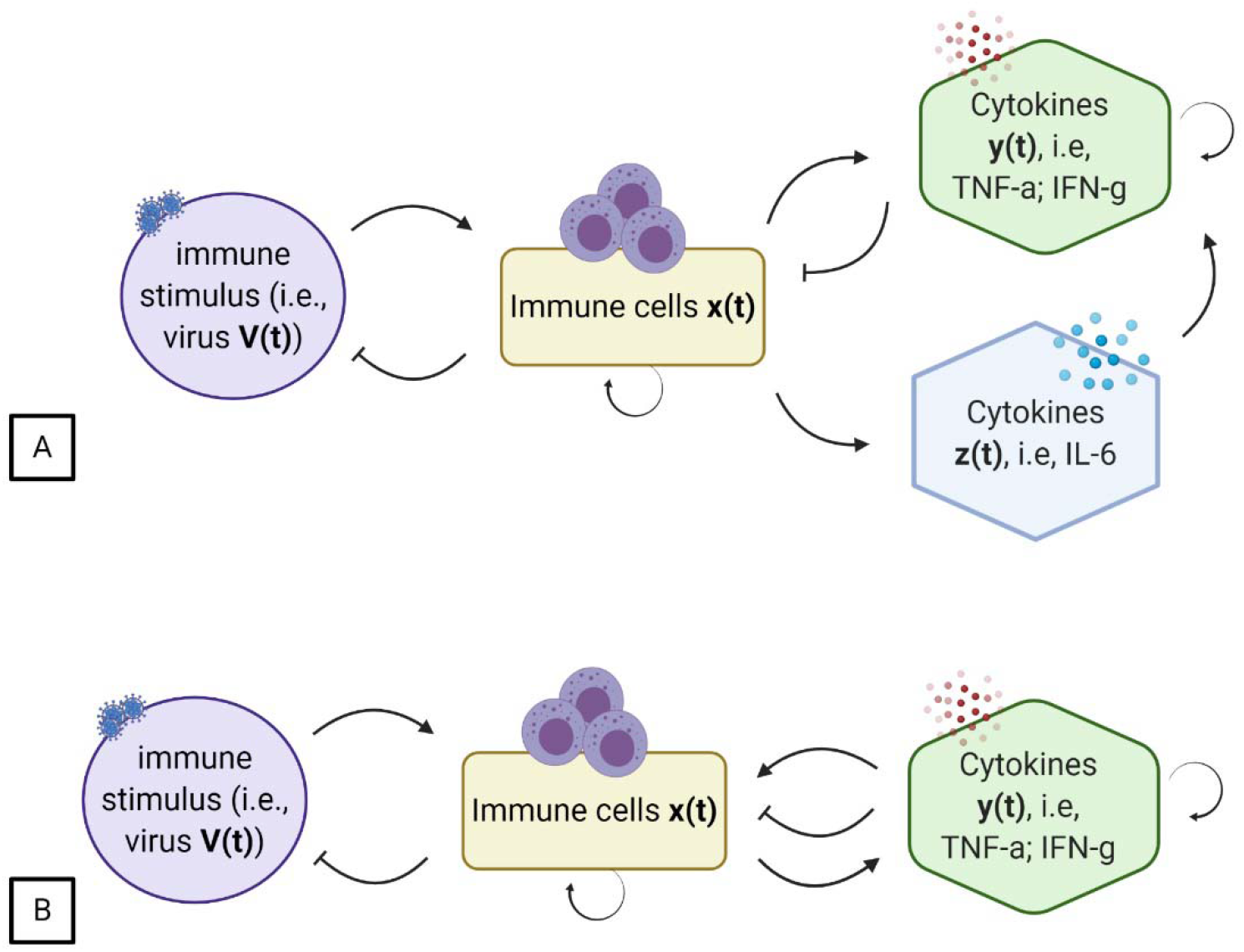
Schematic representation of immune-cytokine interactions subject to perturbation by infection. (A) Full system as described by Equations (1) and (2). (B) Mechanisms described by System (4).

Next, we assume that compared to the dynamics of the immune cells x(t), the y-z subsystem reaches a quasi-steady state before it can affect immune cells x(t).

Therefore, taking 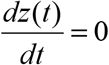 leads to interleukins z(t) reaching a quasi-steady state 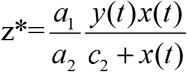. Substituting this expression into System (2), we get the following 2-dimensional system of equations, describing interactions between immune cells and cytokines:

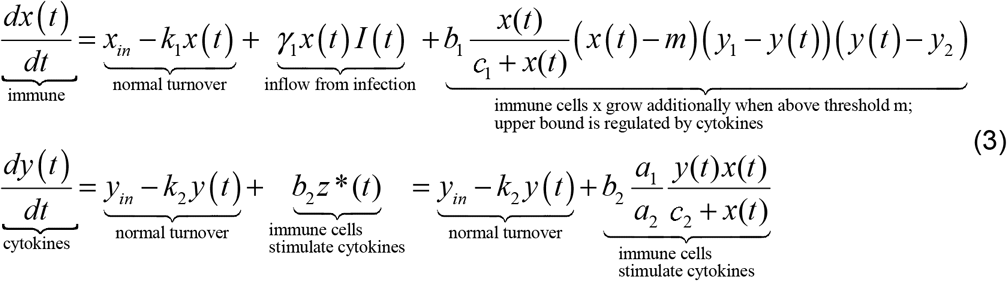

Schematic representation of this reduced system is shown in Figure 1B.

Final system of equations becomes

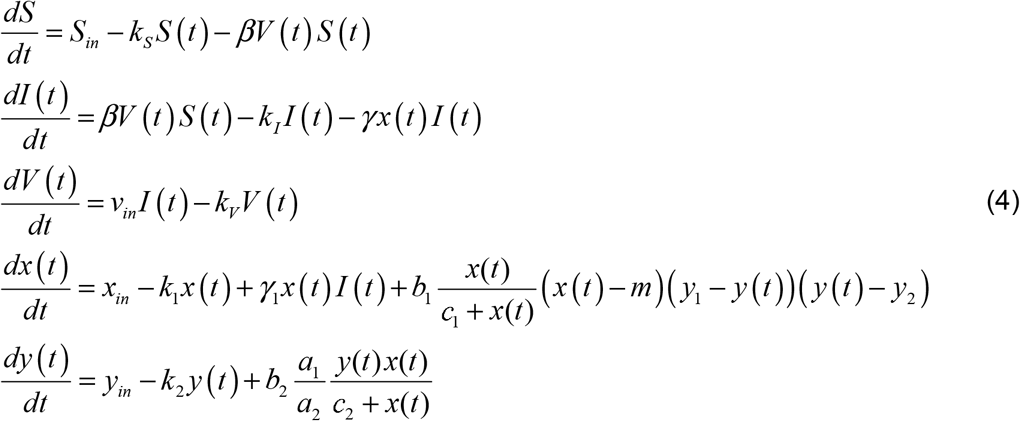

The final System (4) captures the following set of key mechanisms:

1. Viral subsystem serves to provide a stimulus to the immune system that has the potential to trigger cytokine storm in the immune-cytokine subsystem x-y.
2. Immune cells x(t) undergo additional expansion only after threshold m is crossed.
3. Once the threshold m is crossed, cytokines regulate the degree of immune cell expansion as determined by the values of parameters y_1_ and y_2_.

Simulations are conducted as follows. The system is allowed to reach a steady state before infection is introduced at time t=500 (value chosen arbitrarily to ensure sufficient time for the model to reach a steady state). After the infection is introduced, we observe the resulting trajectories of immune cells x(t) and cytokines y(t), as well as the impact of the immune system on the infection.

Due to the phenomenological nature of the proposed model, parameter values were chosen arbitrarily in order to capture qualitatively different behaviors; furthermore, since the model is not fit to specific data, units are chosen to be generic units of volume and time that can be specified when necessary for the purposes of a specific data set. A summary of default parameter values used in the simulations is given in Table 1.

**Table 1.** Parameters used in System (4). Parameter values were chosen arbitrarily to allow to capture qualitatively different behaviors. Parameters a_1_ and a_2_ are taken as 1.

| Parameter | Description | Value | Units |
| --- | --- | --- | --- |
| $S(0)$ | Initial size of population of susceptible cells | 1 | vol. |
| $I(0)$ | Initial size of population of infected cells | 0 | vol. |
| $V(0)$ | Initial size of population of virus particles | 0 | vol. |
| $x(0)$ | Initial size of population of immune cells | 0.07 | vol. |
| $y(0)$ | Initial size of population of cytokines | 0.18 | vol. |
| $S_{in}$ | Production rate of susceptible cells, $S(0) \times k_s$ | 0.01 | vol./time |
| $k_s$ | Normal decay rate of susceptible cells | 0.01 | 1/time |
| $k_I$ | Normal decay rate of infected cells | 0.01 | 1/time |
| $\gamma$ | Rate of elimination of infected cells by immune cells | 0.5 | 1/vol/time |
| $v_{in}$ | Rate of viral replication in infected cells | 0.1 | 1/time |
| $k_v$ | Natural virus decay rate | 0.1 | 1/time |
| $\beta$ | Rate at which virus infects susceptible cells | 0.1 | 1/vol./time |
| $x_{in}$ | Normal production of immune cells, $x(0) \times k_1$ | 7e-4 | vol./time |
| $k_1$ | Normal decay rate of immune cells | 0.01 | 1/time |
| $\gamma_1$ | Conversion of immune cell kill of infected cells into immune cell proliferation | 0.05 | 1/vol/time |
| $m$ | Threshold of activation of additional immune cell proliferation (second touch) | 0.1 | vol. |
| $y_{in}$ | Cytokine production rate, $y(0) \times k_2$ | 0.018 | vol./time |
| $y_1$ | Cytokine-mediated threshold of immune cell expansion | 1 | vol. |
| $y_2$ | Cytokine-mediated threshold of immune cell regulation | 3 | vol. |
| $b_1$ | Rate of additional immune cell expansion as mitigated by cytokines | 1 | 1/(time*vol.<sup>3</sup>) |
| $b_2$ | Rate of cytokine stimulation by immune cells | 1 | 1/time |
| $k_2$ | Normal cytokine decay rate | 0.1 | 1/time |
| $c_1$ | Population size that results in half-maximal growth of $x(t)$ in response to cytokine stimulation | 1 | vol. |
| $c_2$ | Population size that results in half-maximal increase in production of cytokines in response to stimulation by immune cells | 1 | vol. |

## Results

### Dynamical regimes

Initial numerical analysis is performed through variation of parameter b_2_, which represents the impact of immune cells on cytokine production; all other parameters were fixed at values defined in Table 1 unless indicated otherwise.

### Norm

In the first set of simulations we observe expected dynamical behaviors for a normal immune response. Infection at time t=500 is assumed to be sufficiently immunogenic to cause increase in the size of the population of immune cells x(t) for them to surpass threshold m, which now leads to additional immune cell expansion (Figure 2A). As a result, the number of cytokines y(t) increases as well (Figure 2B). Even through for b_2_=0.88, the concentration of y(t) surpasses threshold y_1_, it is not sufficient to initiate additional immune proliferation, and the system quickly returns to equilibrium. The phase-parameter portrait of immune-cytokine interactions is shown in Figure 2C. The immune response is sufficient to clear the infection, as can be seen in Figure 2D.

**Figure 2.**
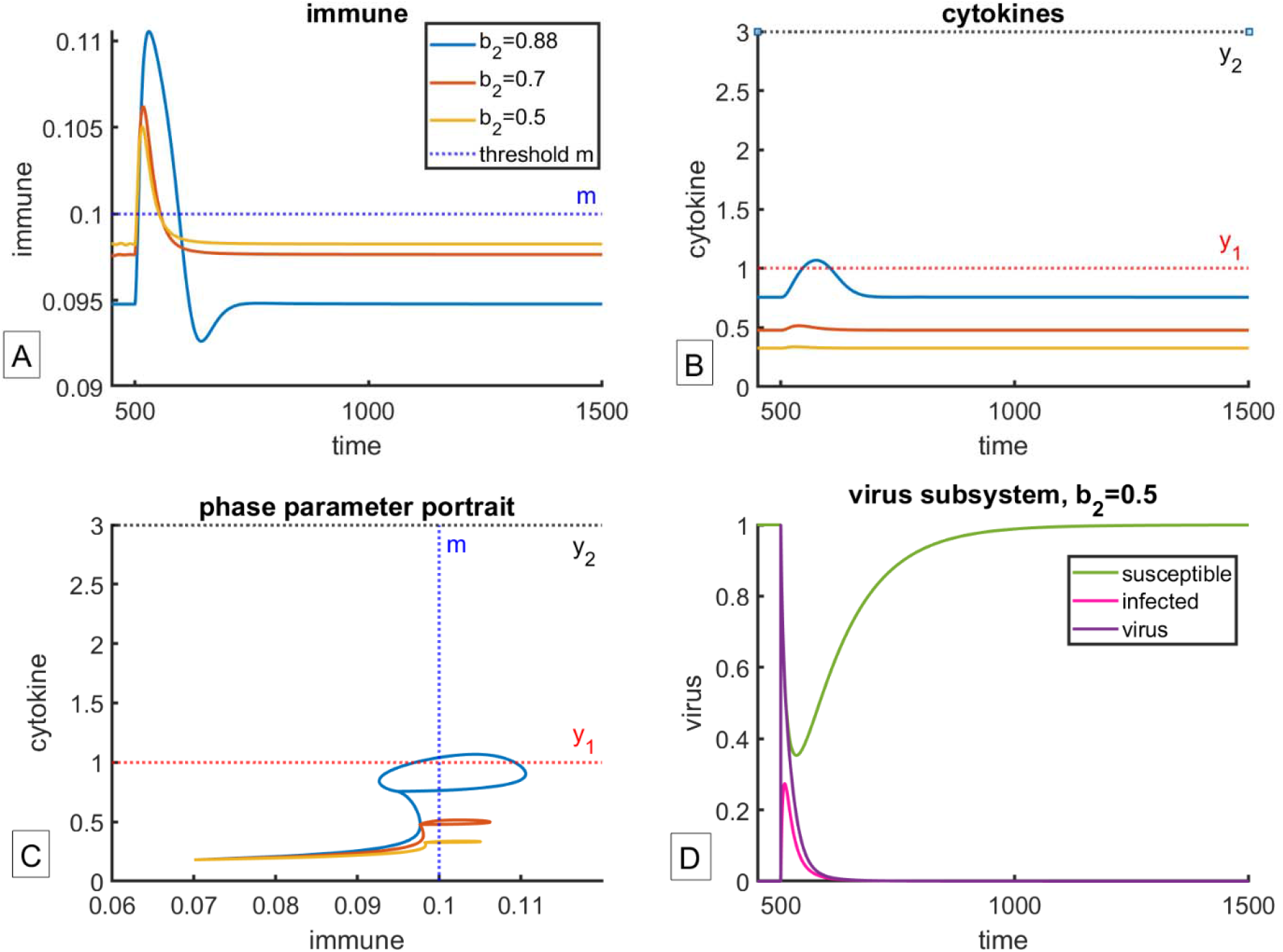
Normal immune response to infection. Infection is introduced at time t=500; parameter b_2_ is increased from 0.5 to 0.7 to 0.88. All other parameters are held constant at values reported in Table 1. (A) Dynamics of immune cells x(t). (B) Dynamics of cytokines y(t). (C) Phase parameter-portrait of the x-y subsystem. (D) Dynamics of the virus subsystem for b_2_=0.5; curves are qualitatively similar for other values of parameter b_2_. After the infection is introduced, the number of susceptible cells decreases, and the number of infected cells increases. This results in increase in immune cells x(t) as population size surpasses threshold m, followed by increase in cytokines y(t). After the infection is cleared, immune cells and cytokines return to pre-infection equilibrium.

Notably, there exists an inverse relationship between baseline levels of immune cells x(t) and cytokines y(t), with lower baseline levels of immune cells corresponding to higher baseline levels of cytokines. While mathematically, this relationship is clearly affected by changes in parameter b_2_, it may also be capturing age-related changes in immune-cytokine balance, with the number of immune cells declining with age, coupled with increased levels of inflammatory cytokines (18). This hypothesis is supported by the observation that older people may be more susceptible to cytokine storms, at least in case of COVID-19 (19).

### Storm

As we increase the value of parameter b_2_, we observe a qualitative change in system behavior, where immune cells and cytokines start amplifying each other, as can be seen in Figure 3 (unless indicated otherwise, in all of the cases shown, the immune system is capable of clearing the virus, and thus the panel with the viral subsystem is not shown). As one can see in Figure 3A, for b_2_=0.89, infection-induced perturbation to the immune system causes a dramatic spike in immune cell population size, leading to subsequent spike in the population of cytokines (Figure 3B), behavior which we interpret as cytokine storm. The phase parameter portrait of the x-y interactions is shown in Figure 3C. While the population eventually returns to equilibrium, it should be noted that after the spike, the model predicts a dip in immune population size before it equilibrates; this prediction remains to be confirmed against experimental observations.

**Figure 3.**
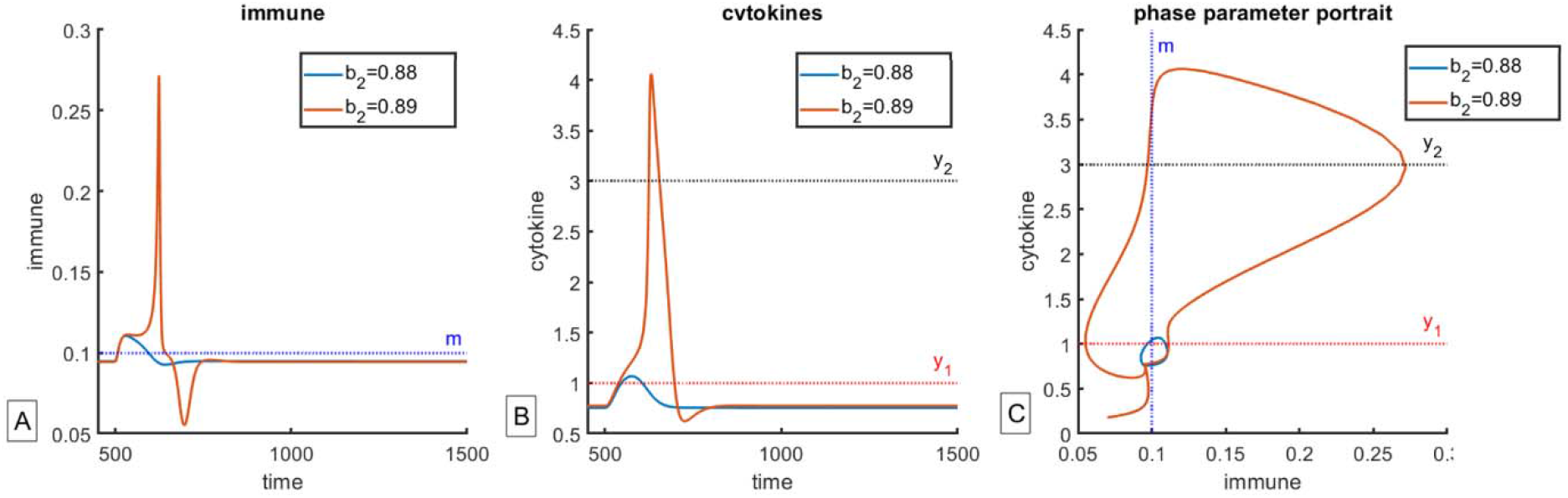
Normal vs storm-like response to infection. As parameter b_2_ increases from 0.88 to 0.89, qualitative change in behavior is observed, as immune cells x(t) and cytokines y(y) start augmenting each other’s behavior. Infection is introduced at time t=500; parameter b_2_ is increased from 0.88 to 0.89. All other parameters are held constant at values reported in Table 1. (A) Dynamics of immune cells x(t). (B) Dynamics of cytokines y(t). (C) Phase parameter-portrait of the x-y subsystem. Dynamics of the virus subsystem is not reported as it is qualitatively similar to one reported in Figure 2D.

### Storms of different magnitude

As we further increase the value of parameter b_2_, we observe that the magnitude of the predicted cytokine storm changes, as does its duration (Figure 4). Moreover, increase in the value of parameter b_2_, which represents the magnitude of cytokine stimulation by the immune cells, results in less severe storms, as can be clearly seen through both the maximal size reached by population of immune cells (Figure 4A), and the size of the characteristic storm-like loop as seen on the phase parameter portrait in Figure 4C.

**Figure 4.**
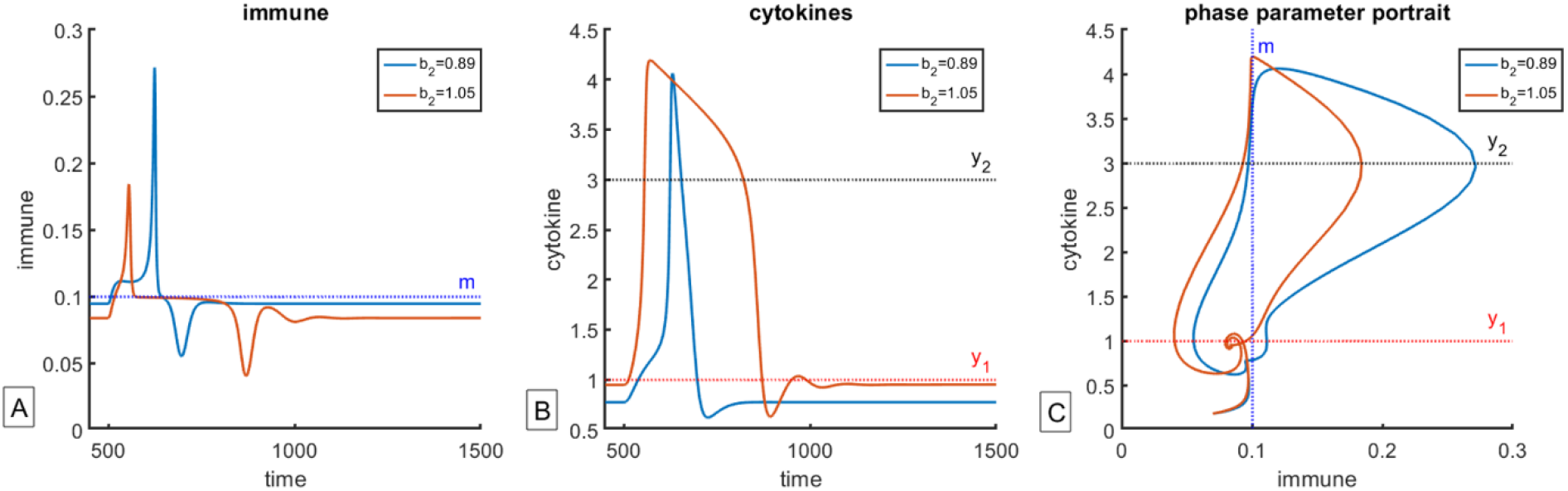
Storms of varying magnitude. As the value of parameter b_2_ increases from 0.89 to 1.05, one can observe storm-like behavior, but the magnitude of the predicted storm is different depending on the value of b_2_. Infection is introduced at time t=500. All other parameters are held constant at values reported in Table 1. (A) Dynamics of immune cells x(t). (B) Dynamics of cytokines y(t). (C) Phase parameter-portrait of the x-y subsystem. Dynamics of the virus subsystem is not reported as it is qualitatively similar to one reported in Figure 2D.

The explanation for this observation lies in timing, and specifically, the amount of time that the population of cytokines y(t) spends between thresholds y_1_ and y_2_ (Figure 5). Larger b_2_ results in increased production of cytokines y(t), and so they reach the inhibitory concentration faster than for smaller values of b_2_, resulting in a shorter and less severe storm-like behavior.

**Figure 5.**
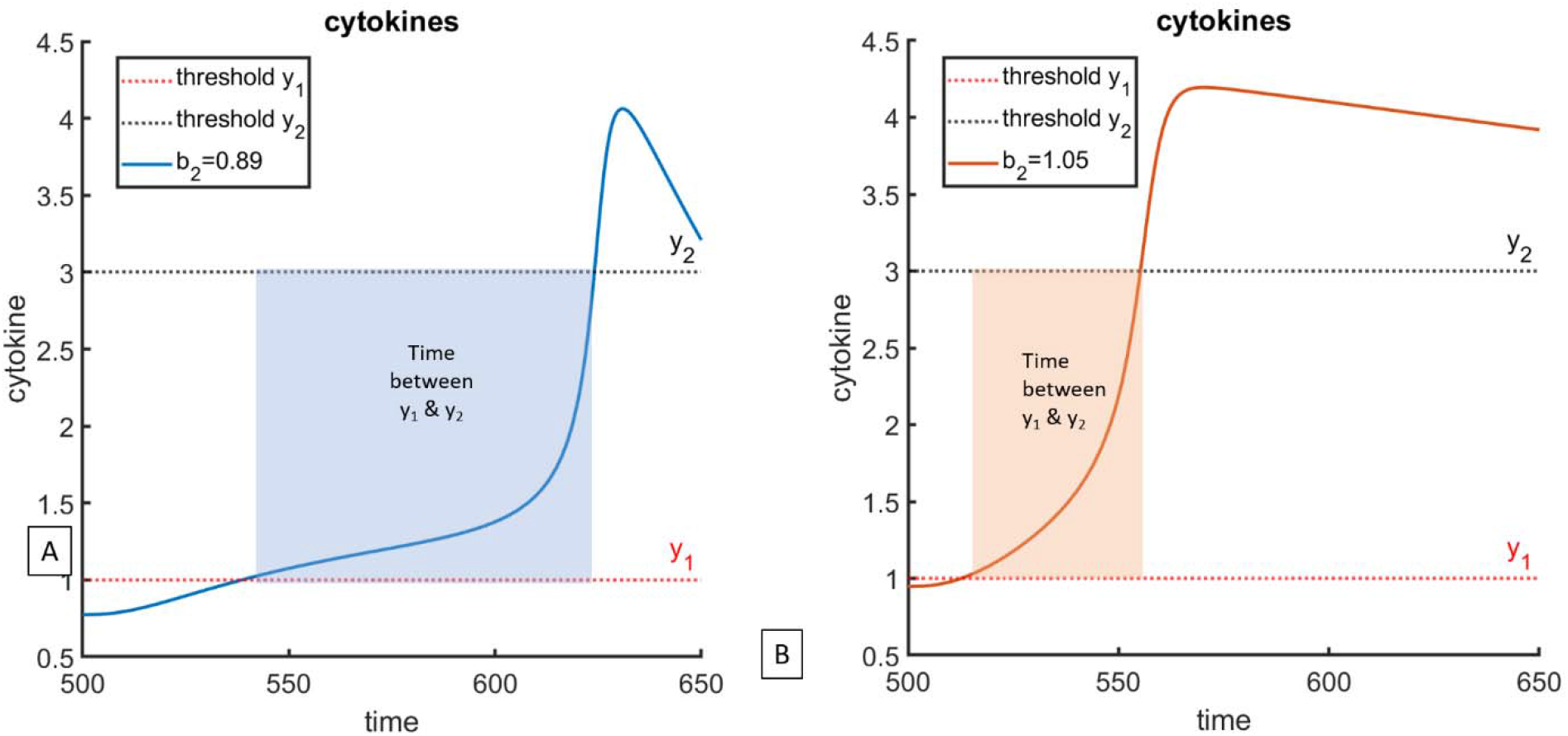
Timing as the key to variations in storm magnitude. (A) Time between thresholds y_1_ and y_2_ for b_2_=0.89. (B) Time between thresholds y_1_ and y_2_ for b_2_=1.05. Since b_2_ represents stimulation of cytokines by the immune cells, larger values of b_2_ result in faster time between thresholds y_1_ and y_2_, resulting in a storm of a smaller magnitude.

### New norm

Finally, as we further increase the value of parameter b_2_, we observe the population reaching a new equilibrium, with population of immune cells x(t) equilibrating at the threshold m, which is higher than pre-disease baseline; in this case, cytokines equilibrate above threshold y_2_ (Figure 6). We propose that this behavior can be interpreted as infection-induced autoimmunity, a phenomenon that has been previously reported in the literature (20).

**Figure 6.**
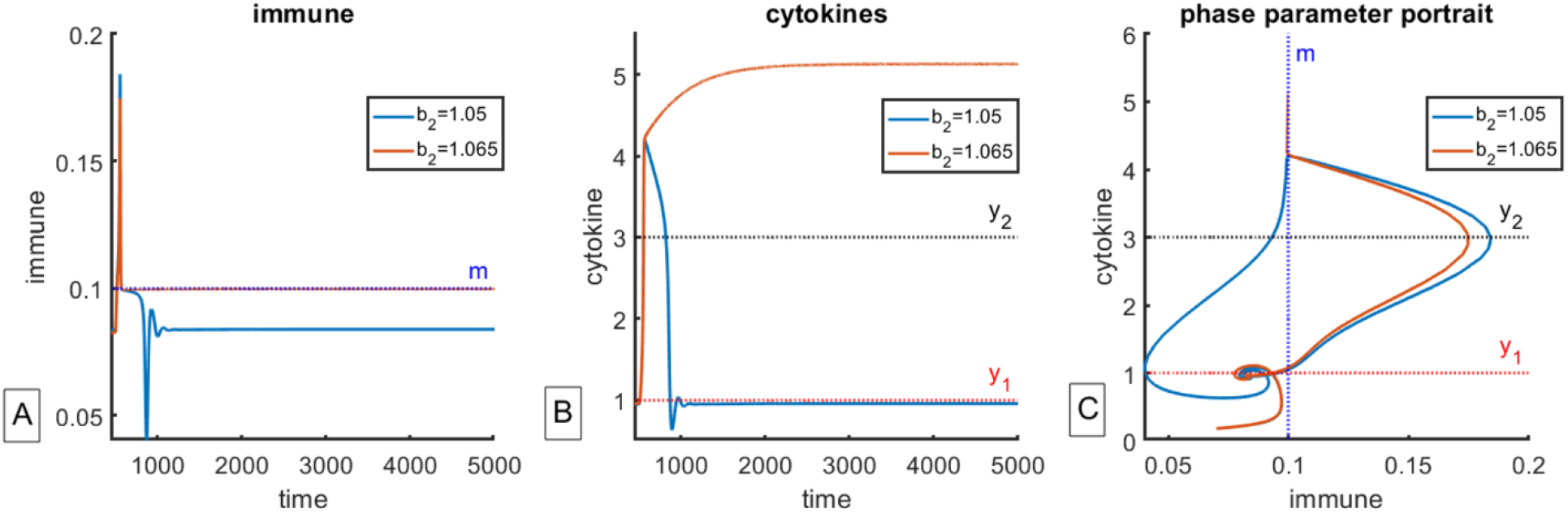
New norm. As the value of parameter b_2_ increases from 1.05 to 1.065, one can observe shift towards a new equilibrium, where immune cells x(t) equilibrate at threshold m, and cytokines y(t) equilibrate above threshold y_2_. Infection is introduced at time t=500. All other parameters are held constant at values reported in Table 1. (A) Dynamics of immune cells x(t). (B) Dynamics of cytokines y(t). (C) Phase parameter-portrait of the x-y subsystem. Dynamics of the virus subsystem is not reported as it is qualitatively similar to one reported in Figure 2D.

### Sequence of dynamical regimes

Next, we wanted to capture the impact on system dynamics of variation of parameter b_1_, which represents the rate at which cytokines y(t) stimulate immune system x(t); all other parameters were held constant at values given in Table 1. The result is shown in Figure 7, which reveals a sequence of dynamical regimes, where cytokine storm is a transient regime that can become realized when several conditions are met. Specifically, we have shown that for low b_1_, the immune response is insufficient to clear the infection (region 0), regardless of the value of b_2_. Once the value of b_1_ is sufficiently large, we can observe that increasing b_2_ leads first to normal response (region 1), after which the immune system quickly returns to pre-disease equilibrium. As we increase b_2_, we observe storm-like behavior, with smaller b_2_ predicting more severe storms due to longer time spent between thresholds y_1_ and y_2_ (region 2). Further increase in b_2_ leads to less severe storms because of shorter time spent between y_1_ and y_2_ (region 3). Finally, further increase of b_2_ results in what we term a “new norm”, or infection-induced autoimmunity (region 4).

**Figure 7.**
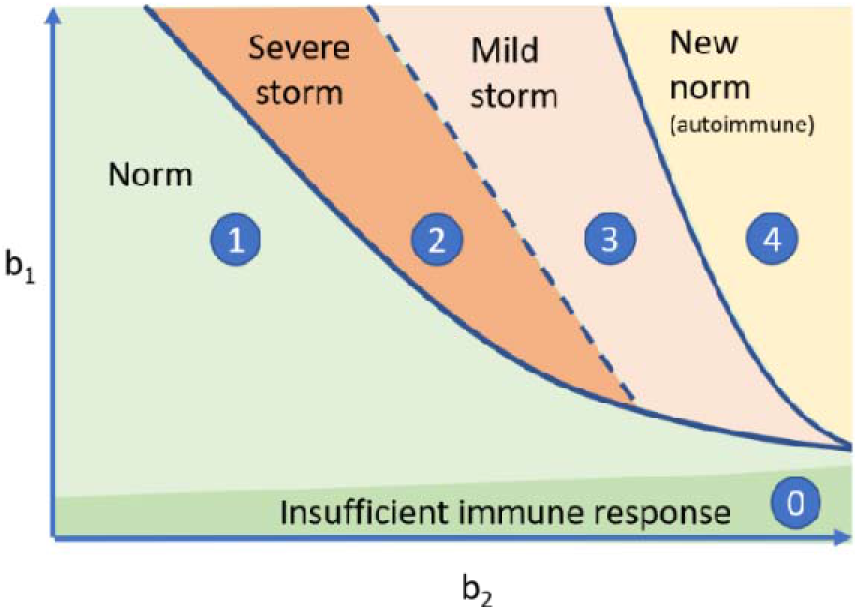
Sequence of regimes predicted by the model, subject to variation of parameters b_1_ and b_2_, where cytokine storm is revealed to be a transient regime.

### Conditions corresponding to storm-like behavior

Additional insights into observed behaviors can be obtained from analysis of isoclines and the change in their relative positions depending on values of parameters within the relevant parameter space; parameter values are held at values reported in Table 2 unless indicated otherwise. Recall that we are only considering the case when stable disease-free equilibrium is such that x*<m, and additional immune cell expansion only occurs after this threshold is passed as a result of perturbation, either from infection or any other cause.

**Table 2.**
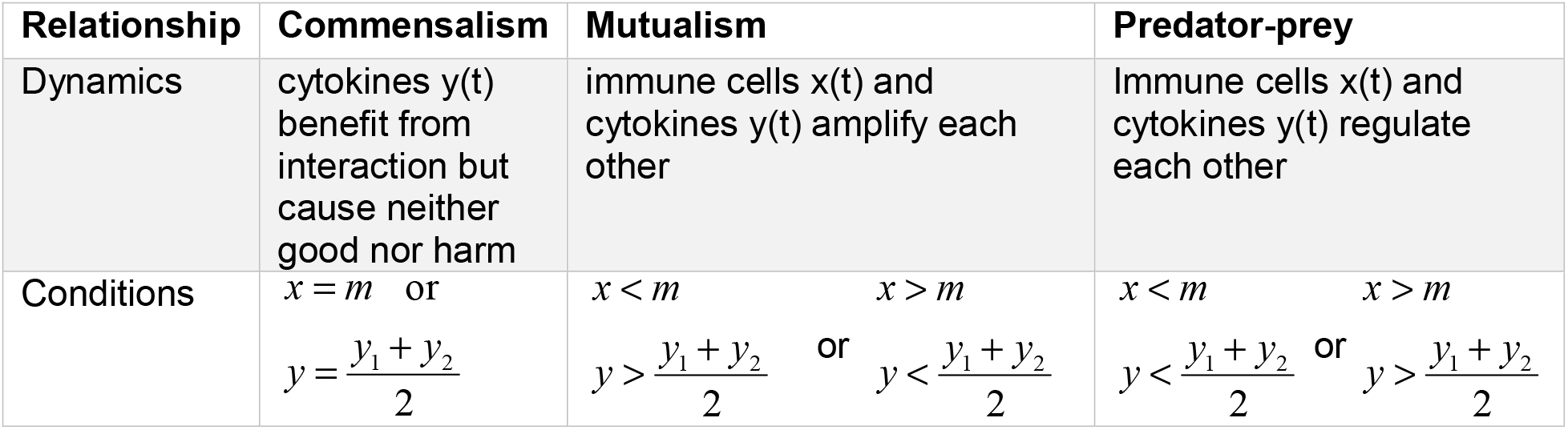
Ecological relationships between immune cells x(t) and cytokines y(t).

Isoclines for System (3) are given by

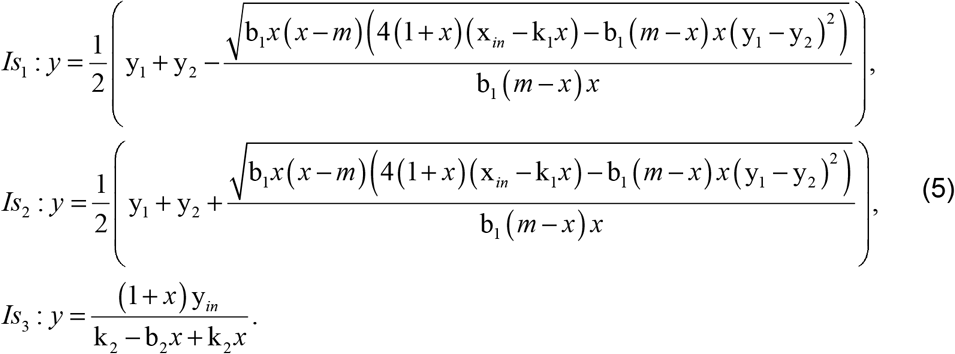

Depending on parameter values, isoclines can have between one and three points of intersection. As one can see in Figure 8, there always exists one stable equilibrium, a nodal sink, which corresponds to infection-free immune-cytokine balance. Additionally, there can exist two more equilibrium points, a spiral source and a saddle point, which exist for small values of b_2_ (Figure 8A); as b_2_ increases, the source and the saddle merge (Figure 8B) and eventually disappear (Figure 8C), resulting in existence only of the nodal sink.

**Figure 8.**
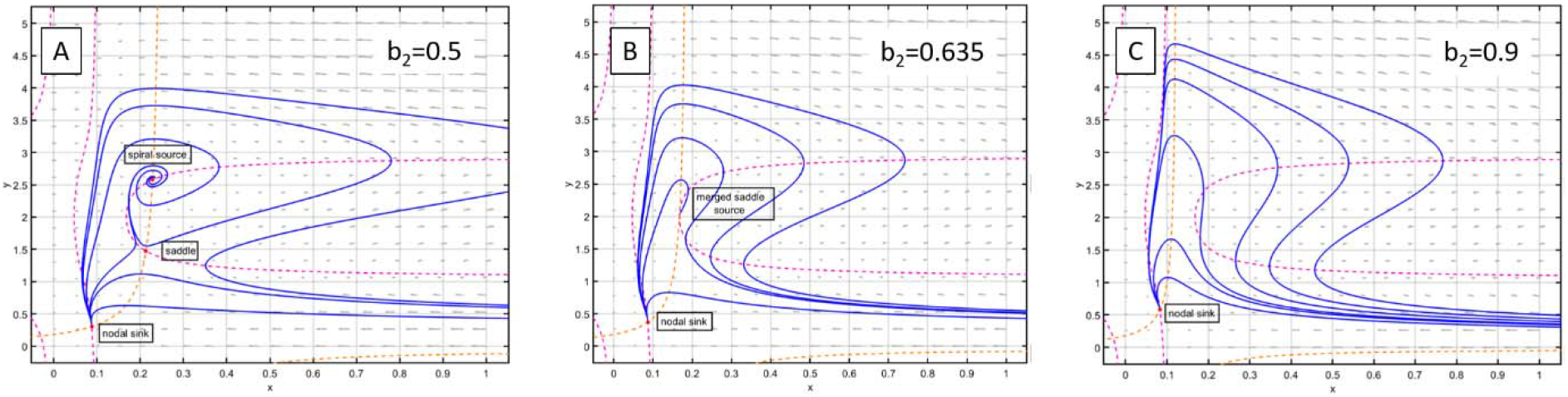
Isocline analysis of immune-cytokine subsystem (3). (A) For smaller b_2_, there can exist 3 equilibrium points, one stable node, one spiral source and a saddle point. (B) As the value of b_2_ increases, saddle and source merge into a single point. (C) As b_2_ increases further, only one equilibrium point remains. We observe that storm-like dynamics occurs only when there exists a single equilibrium point.

In this system, we observed that storm-like dynamics occur only when there exists only one equilibrium point (Figure 8C).

#### Ecological perspective

To further our understanding of this system, we analyze it from the perspective of community modules, which are frequently used in ecological systems (21). Consider partial derivatives of immune-cytokine subsystem (3):

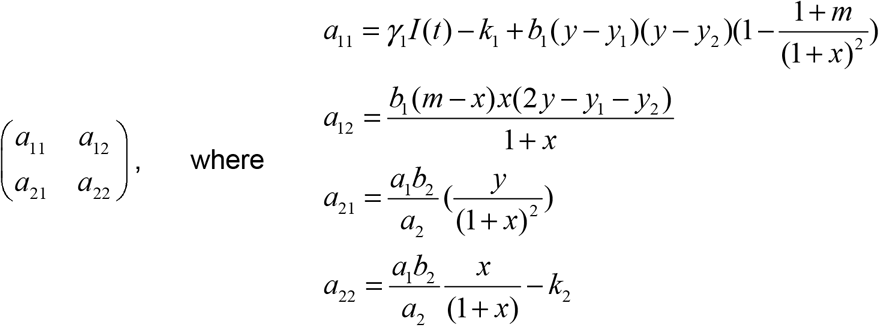

Recall from (21) that if a_21_ is > 0, depending on the sign of a_12_, the relationship between the two variables can be either mutualistic if a_21_>0, or predator-prey if a_12_<0. In a mutualistic system, 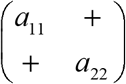, populations amplify each other, while in a predator-prey type system, 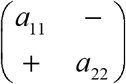, the two interacting populations regulate each other. Within the context of the proposed immune-cytokine System (3), one can classify observed dynamical regimes depending on the sign of a_12_ as follows.

The two populations are in a mutualistic relationship when 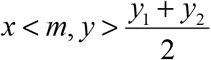 or when 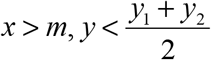; in this case, immune cells x(t) and cytokines y(t) amplify each other, which corresponds to regions of accelerated immune and cytokine population size increase as observed in Figure 9. The two populations are in a predator-prey type relationship if 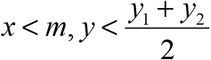 or 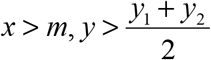; in this case cytokines act as regulators and “dampeners” of immune response. Notably, if a_12_=0, then the two populations are in a commensal relationship, where cytokines y(t) benefit from the interactions but cause neither increase nor decrease to the immune population size. This occurs when *x* = *m* or 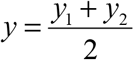, a behavior we observe in the “new norm” region of Figure 7.

**Figure 9.**
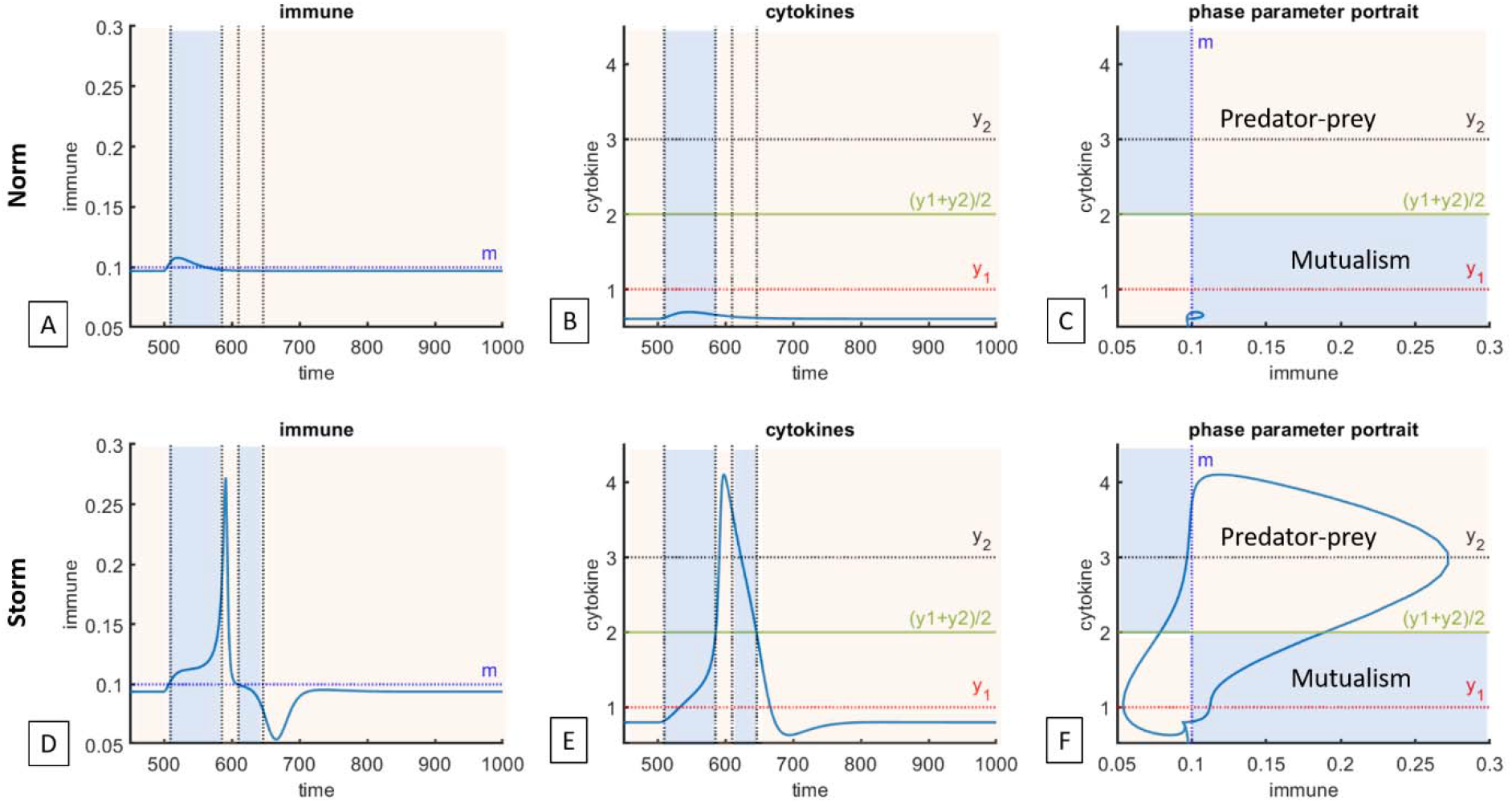
Application of ecological analysis to immune-cytokine trajectories for normal and storm-like responses. Boundaries for predator-prey vs mutualism interactions are given in Table 2. Top panel: norm, b_2_=0.8, other parameters reported in Table 1. Dashed lines correspond to conditions when switch from mutualism to predator-prey like behavior can occur. (A) Immune cells x(t); (B) cytokines y(t); (C): phase parameter portrait. Normal immune response involves a single transition from stabilizing predator-prey type interaction to mutually amplifying mutualist and back to stabilizing predator-prey. Bottom panel: cytokine storm, b_2_=0.9. (D) immune cells x(t), (E) cytokines y(t), (F) phase-parameter portrait. In a cytokine storm, there exists an additional predator-prey to mutualism cycle compared to normal response.

These results are summarized in Table 2 and visualized in Figure 9.

Notably, this perspective could provide potential additional explanation for why timing matters in treatment administration: if a cytokine blocker results in reducing cytokine concentration such that the system moves into, or remains in a mutualistic regime, then it may instead amplify the severity of immune and cytokine production rather than reduce its impact.

### Impact of parameter γ and the severity of infection

Up to this point, we have identified the impact of the following parameters on occurrence of a cytokine storm: 1) parameters b_1_ and b_2_, which represent the degree to which immune cells and cytokines stimulate each other’s production, 2) parameter m, which represents a threshold for additional immune cell expansion, and 3) parameters y_1_ and y_2_, which determine a region of cytokine-induced stimulation or inhibition of additional immune cell expansion.

Now we evaluate the impact of responsiveness of immune system to infected cells themselves as measured through changes in the value of parameter *γ*. We fix the value of b_1_ and vary parameters b_2_ and *γ* to evaluate whether the infection was cleared, and whether the immune-cytokine response is normal or storm-like. As one can see in schematic Figure 10, the model predicts that for large enough values of *γ*, the infection will be cleared without a cytokine storm; it also confirms that increase in the value of b_2_ can lead to storm-like behavior. Notably, the model also predicts the possibility of a cytokine storm without infection clearance (figure not shown, parameter values are b_2_=0.9, *γ* = 0.1, b_1_=1; other parameters are as reported in Table 1). In this case, the immune system is not efficient in clearning the infection (small *γ*) but the cytokine-immune dynamics are triggered, resulting storm-like dynamics due to a combination of individual-specific intrinsic factors summarized above.

**Figure 10.**
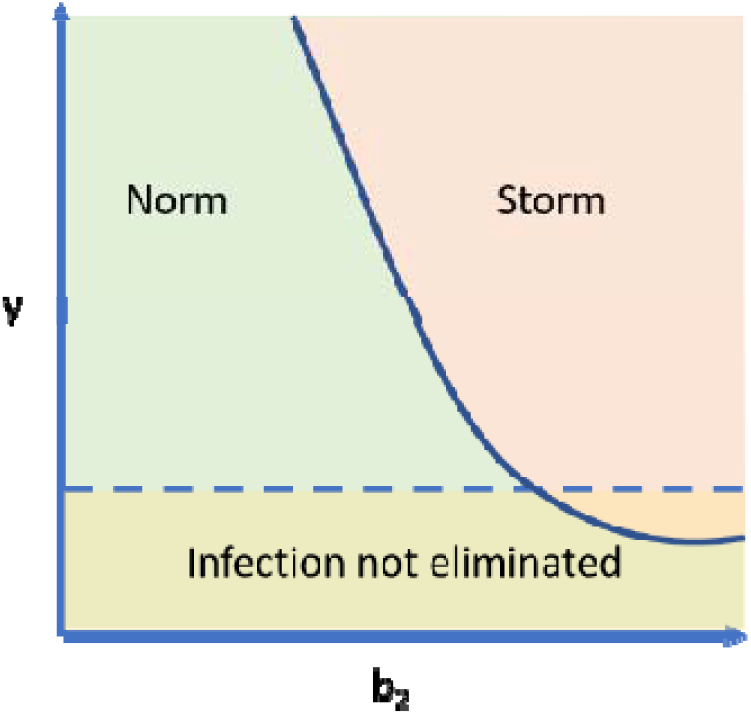
Impact of variation of immungenicity parameter *γ* on immune response. It is possible to observe both normal and storm-like reponse, with or without infection elimination.

### Chronic infection and the long-haulers

Long-haulers are a subset of patients who develop chronic coronavirus disease (22–24). Within the frameworks of the proposed model, this behavior is captured as stable non-trivial equilibrium between all five variables of System (4), as can be seen in Figure 11. Notably, as predicted by analysis done in Figure 10, this can occur with or without storm-like dynamics.

**Figure 11.**
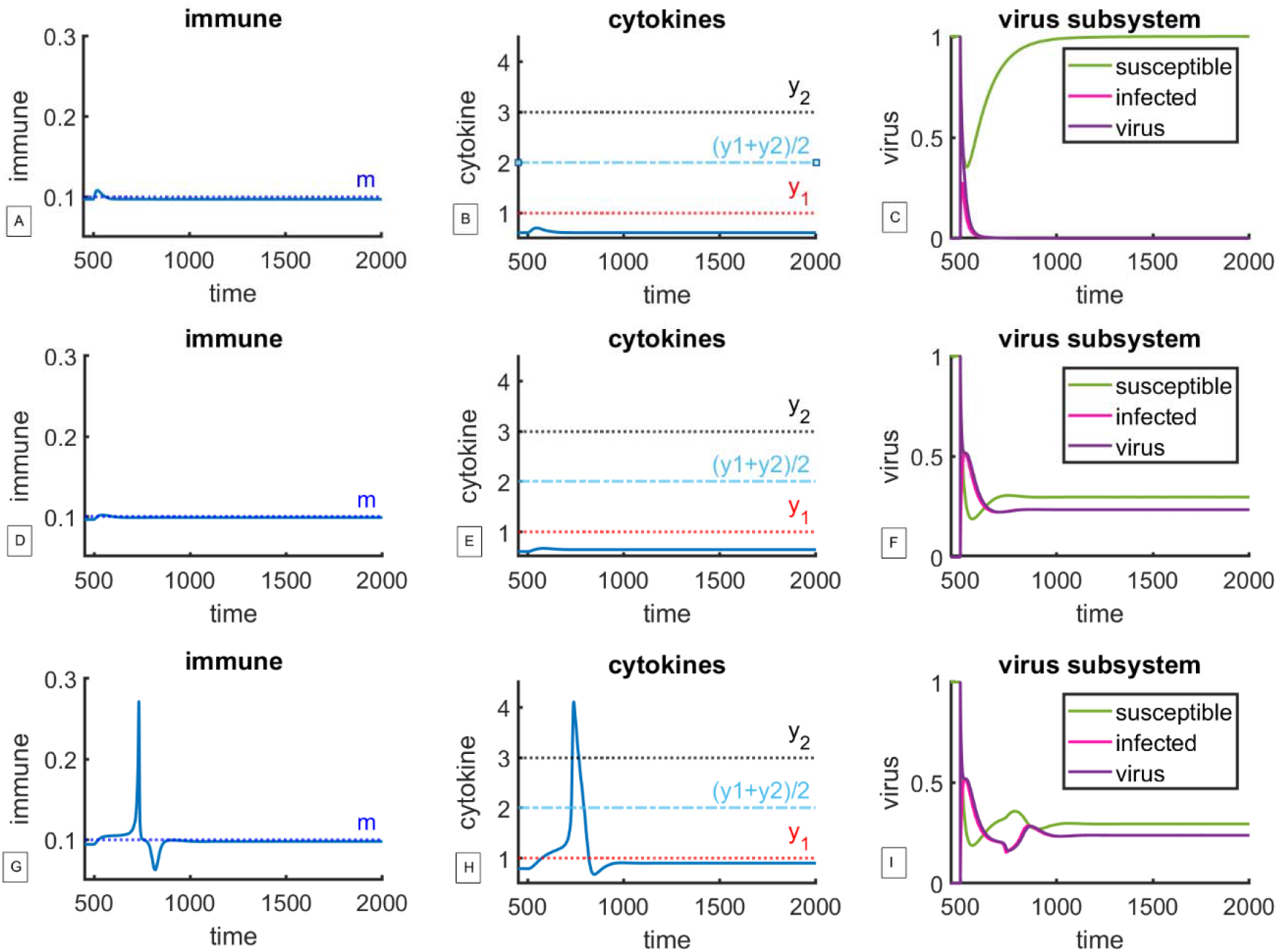
Model predicts possibility of chronic infection (long haulers) with and without cytokine storm. Top panel: normal immune response, *γ* =0.5, b_2_=0.8, other parameters reported in Table 1. (A) Immune cells x(t), (B) cytokines y(t), (C) viral subsystem. Infection is eliminated. Middle panel, normal immune response; *γ* =0.1, b_2_=0.8. (D) immune cells x(t), (E) cytokines y(t), (F) virus subsystem. Even though immune-cytokine dynamics are normal, the efficiency of infection kill is too low, resulting in persistent infection, which can be interpreted as a “long hauler”. Bottom panel: b_2_=0.9, *γ* =0.1. (G) immune cells x(t), (H) cytokines y(t), (I) viral subsystem. Even though immune-cytokine dynamics show cytokine storm, the efficiency of infection elimination is insufficient, suggesting that an individual can go through a cytokine storm and still not clear the infection. Note: in figures A, D and G, immune system equilibrates below threshold m, returning to its pre-disease baseline. The y-axis was scaled to enable comparison between the cases.

## Discussion

Here we propose a conceptual mathematical model of immune-cytokine interactions capable of reproducing the qualitative behaviors that capture transition from normal immune response to a response that can be interpreted as cytokine storm. The goal of the model was not to describe a particular data set or to incorporate great biological detail but to capture qualitative relationships between the broad classes of immune cells and cytokines that are sufficient to reproduce these dynamics, as well as to identify key parameters that may suggest whether an individual may be susceptible to experiencing a cytokine storm. The proposed model was coupled with a SIV model that describes immune response to a viral infection and which serves to trigger immune-cytokine interactions. The viral subsystem serves as a source of perturbation and is not the focus of the current discussion; it was chosen nevertheless to enable demonstration of various dynamical regimes, such as chronic infection, and can be substituted by another model tailored to the question of interest.

We show that there exists a parameter-dependent sequence of dynamical regimes (Figure 7) that describe how immune cells and cytokines stimulate each other in response to infection as the body tries to mount an appropriately strong immune response while also avoiding excessive activation. Specifically, we show that as the value of parameter b_2_ (extent of cytokine stimulation by immune cells) increases, we see a transition from normal response (Figures 2 and 3) to cytokine storm (Figure 4) to a regime that we interpret as infection-induced autoimmunity (Figure 6). We also demonstrate that counterintuitively, lower b_2_ predicts more severe storm-like behavior due to longer time spent between cytokine-specific thresholds y_1_ and y_2_ (Figure 5). If the framework proposed here is true, then susceptibility to a cytokine storm is more likely to be an individual-specific characteristic that may or may not become realized subject to a challenge to the immune system. The model also predicts the existence of so-called long-haulers, patients harboring a chronic infection that may or may not be accompanied by storm-like immune-cytokine dynamics (Figure 11).

The proposed immune-cytokine model is reduced to two equations, which allows for additional analysis. Specifically, a 2-dimensional system was analyzed from the point of view of ecological community modules, revealing conditions under which the immune cells and cytokines were in a mutually amplifying mutualistic vs more stabilizing predator-prey type relationship (Table 2). We were able to show the difference between normal and storm-like behavior from the point of view of switching between the two types of ecological relationships (Figure 9), where an additional mutualistic phase amplifies storm-like behavior.

Through our analysis, we demonstrate that within the frameworks of the proposed model, cytokine storm is a transient regime that can become realized when the following individual and infection-specific conditions are met:

1. when baseline level of immune cells is close to activation threshold m,
2. when cytokines spend a lot of time between thresholds y_1_ and y_2_, either because the two thresholds are far apart, or when the value of parameter b_2_ is small, and
3. when the infection is sufficiently immunogenic.

Even through here the perturbation to immune-cytokine equilibrium was achieved using a viral subsystem, other model variations can be used in future work, including simulations of impact of therapeutic agents that are known to have a high likelihood of cytokine storm reaction, such as bispecific T cell engagers (BiTEs) or CAR-T cell therapies (5,6,25,26). Furthermore, since two of the three identified factors that can result in a storm-like reaction to an immunological challenge are individual-specific, it is likely that they can be leveraged during patient selection process for such therapies if a sufficiently robust approach to estimating these qualities can be found, such as genetic factors that may serve as predictive biomarkers (27). It is our hope that the proposed model can help narrow down the list of possible culprits responsible for cytokine storms and guide additional research into ways that it can be mitigated.

## Acknowledgements

The authors would like to thank Fred Adler and Joel Brown for looking over the earlier drafts of the model and providing valuable comments and insights.

## References

1. Ye Q, Wang B, Mao J. The pathogenesis and treatment of theCytokine Storm’in COVID-19. Journal of infection. Elsevier; 2020;80(6):607–613.

2. Ferrara JL. Cytokine dysregulation as a mechanism of graft versus host disease. Current opinion in immunology. Elsevier; 1993;5(5):794–799.

3. Huang K-J, Su I-J, Theron M, Wu Y-C, Lai S-K, Liu C-C, et al. An interferon-γ-related cytokine storm in SARS patients. Journal of medical virology. Wiley Online Library; 2005;75(2):185–194.

4. Oxford JS, Gill D. Unanswered questions about the 1918 influenza pandemic: origin, pathology, and the virus itself. The Lancet Infectious Diseases. Elsevier; 2018;18(11):e348–e354.

5. Porter D, Frey N, Wood PA, Weng Y, Grupp SA. Grading of cytokine release syndrome associated with the CAR T cell therapy tisagenlecleucel. Journal of hematology & oncology. Springer; 2018;11(1):35.

6. Chen X, Kamperschroer C, Wong G, Xuan D. A Modeling Framework to Characterize Cytokine Release upon T-Cell–Engaging Bispecific Antibody Treatment: Methodology and Opportunities. Clinical and Translational Science. Wiley Online Library; 2019;12(6):600–608.

7. Suntharalingam G, Perry MR, Ward S, Brett SJ, Castello-Cortes A, Brunner MD, et al. Cytokine storm in a phase 1 trial of the anti-CD28 monoclonal antibody TGN1412. New England Journal of Medicine. Mass Medical Soc; 2006;355(10):1018–1028.

8. Fajgenbaum DC, June CH. Cytokine storm. New England Journal of Medicine. Mass Medical Soc; 2020;383(23):2255–2273.

9. Tisoncik JR, Korth MJ, Simmons CP, Farrar J, Martin TR, Katze MG. Into the eye of the cytokine storm. Microbiology and Molecular Biology Reviews. Am Soc Microbiol; 2012;76(1):16–32.

10. Waito M, Walsh SR, Rasiuk A, Bridle BW, Willms AR. A mathematical model of cytokine dynamics during a cytokine storm. Mathematical and Computational Approaches in Advancing Modern Science and Engineering. Springer; 2016. p. 331–339.

11. Yiu HH, Graham AL, Stengel RF. Dynamics of a cytokine storm. PloS one. Public Library of Science; 2012;7(10):e45027.

12. Hopkins B, Tucker M, Pan Y, Fang N, Huang ZJ. A model-based investigation of cytokine storm for T-cell therapy. IFAC-PapersOnLine. Elsevier; 2018;51(19):76–79.

13. Baker M, Denman-Johnson S, Brook BS, Gaywood I, Owen MR. Mathematical modelling of cytokine-mediated inflammation in rheumatoid arthritis. Mathematical medicine and biology: a journal of the IMA. OUP; 2013;30(4):311–337.

14. Zhang W, Jang S, Jonsson CB, Allen LJ. Models of cytokine dynamics in the inflammatory response of viral zoonotic infectious diseases. Mathematical medicine and biology: a journal of the IMA. Oxford University Press; 2019;36(3):269–295.

15. Du SQ, Yuan W. Mathematical modeling of interaction between innate and adaptive immune responses in COVID-19 and implications for viral pathogenesis. Journal of Medical Virology. Wiley Online Library; 2020;

16. Karki R, Sharma BR, Tuladhar S, Williams EP, Zalduondo L, Samir P, et al. COVID-19 cytokines and the hyperactive immune response: Synergism of TNF-α and IFN-γ in triggering inflammation, tissue damage, and death. bioRxiv. Cold Spring Harbor Laboratory; 2020;

17. Ley K. The second touch hypothesis: T cell activation, homing and polarization. F1000Research. Faculty of 1000 Ltd; 2014;3.

18. Montecino-Rodriguez E, Berent-Maoz B, Dorshkind K, others. Causes, consequences, and reversal of immune system aging. The Journal of clinical investigation. Am Soc Clin Investig; 2013;123(3):958–965.

19. Mueller AL, McNamara MS, Sinclair DA. Why does COVID-19 disproportionately affect older people? Aging. 2020;12(10).

20. Kivity S, Agmon-Levin N, Blank M, Shoenfeld Y. Infections and autoimmunity–friends or foes? Trends in immunology. Elsevier; 2009;30(8):409–414.

21. Holyoak M, Leibold MA, Holt RD. Metacommunities: spatial dynamics and ecological communities. University of Chicago Press; 2005.

22. Siegelman JN. Reflections of a COVID-19 Long Hauler. JAMA. American Medical Association; 2020;324(20):2031–2032.

23. Carfi A, Bernabei R, Landi F, others. Persistent symptoms in patients after acute COVID-19. Jama. American Medical Association; 2020;324(6):603–605.

24. Baig AM. Chronic COVID Syndrome: Need for an appropriate medical terminology for Long-COVID and COVID Long-Haulers. Journal of medical virology. Wiley Online Library; 2020;

25. Bonifant CL, Jackson HJ, Brentjens RJ, Curran KJ. Toxicity and management in CAR T-cell therapy. Molecular Therapy-Oncolytics. Elsevier; 2016;3:16011.

26. Hosseini I, Gadkar K, Stefanich E, Li C-C, Sun LL, Chu Y-W, et al. Mitigating the risk of cytokine release syndrome in a Phase I trial of CD20/CD3 bispecific antibody mosunetuzumab in NHL: impact of translational system modeling. NPJ systems biology and applications. Nature Publishing Group; 2020;6(1):1–11.

27. Wurfel M. Genetic insights into sepsis: what have we learned and how will it help? Current pharmaceutical design. Bentham Science Publishers; 2008;14(19):1900–1911.

